# Phosphorylation of the ancestral histone variant H3.3 amplifies stimulation-induced transcription

**DOI:** 10.1101/808048

**Authors:** Anja Armache, Shuang Yang, Lexi E Robbins, Ceyda Durmaz, Andrew W Daman, Jin Q Jeong, Alexia Martínez de Paz, Arjun Ravishankar, Tanja Arslan, Shu Lin, Tanya Panchenko, Benjamin A. Garcia, Sandra B. Hake, Haitao Li, C. David Allis, Steven Z. Josefowicz

## Abstract

Complex organisms are able to rapidly induce select genes among thousands in response to diverse environmental cues. This occurs in the context of large genomes condensed with histone proteins into chromatin. The macrophage response to pathogen sensing, for example, rapidly engages highly conserved signaling pathways and transcription factors (TFs) for coordination of inflammatory gene induction^1–3^. Enriched integration of histone H3.3, the ancestral histone H3 variant, is a feature of inflammatory genes and, in general, dynamically regulated chromatin and transcription^4–7^. However, little is known of how chromatin is regulated at rapidly induced genes and what features of H3.3, conserved from yeast to human, might enable rapid and high-level transcription. The amino-terminus of H3.3 contains a unique serine residue as compared with alanine residues found in “canonical” H3.1/2. We find that this H3.3-specific serine residue, H3.3S31, is phosphorylated (H3.3S31ph) in a stimulation-dependent manner along the gene bodies of rapidly induced response genes in mouse macrophages responding to pathogen sensing. Further, this selective mark of stimulation-responsive genes directly engages histone methyltransferase (HMT) SETD2, a component of the active transcription machinery. Our structure-function studies reveal that a conserved positively charged cleft in SETD2 contacts H3.3S31ph and specifies preferential methylation of H3.3S31ph nucleosomes. We propose that features of H3.3 at stimulation induced genes, including H3.3S31ph, afford preferential access to the transcription apparatus. Our results provide insight into the function of ancestral histone variant H3.3 and the dedicated epigenetic mechanisms that enable rapid gene induction, with implications for understanding and treating inflammation.

---

A poorly understood feature of stimulation-induced genes is their ability to effectively engage the general transcription machinery for rapid expression. Selective, induced gene transcription, for example during heat shock^8^ or the inflammatory response, occurs rapidly and robustly, despite these genes’ *de novo* expression among thousands of constitutively expressed genes. We considered that stimulation-induced transcription may be controlled by dedicated epigenetic mechanisms in cooperation with signal-activated transcription factors (TFs). Among stimulation-responsive features of chromatin, histone phosphorylation can be an efficient and potent means of transmitting signals via kinase cascades to chromatin regions associated with stimulation-responsive genes with the potential to augment their transcription^9–14^.

H3.3 is the conserved, ancestral H3 variant and the only H3 present in some simple eukaryotes, including *S. cerevisiae*. In complex organisms, H3.3 is uniquely expressed outside of the cell cycle and plays a variety of roles in transcription, genomic stability and mitosis, while so-called “canonical” H3.1/2 histones are expressed in a “replication-dependent” manner and provide a principal packaging role to accommodate the doubling genome^15,16^. The amino-terminal H3.3 ‘tail’ differs from that of H3.1/2 by a single amino acid, a serine at position 31 in H3.3 in place of an alanine in H3.1/2 (Fig. 1A and fig. S1A). Despite the well-characterized enrichment of H3.3 in dynamic chromatin, the potential regulatory roles of H3.3S31 and H3.3-specific phosphorylation are unknown^4–7,17^. Here, we report that H3.3 phosphorylation at the conserved and H3.3-specific serine 31 (H3.3S31ph) amplifies the rapid, high-level transcription of stimulation-induced gene expression. We present a specific biophysical mechanism that provides these select genes with augmented transcriptional capacity.

**Figure 1:**
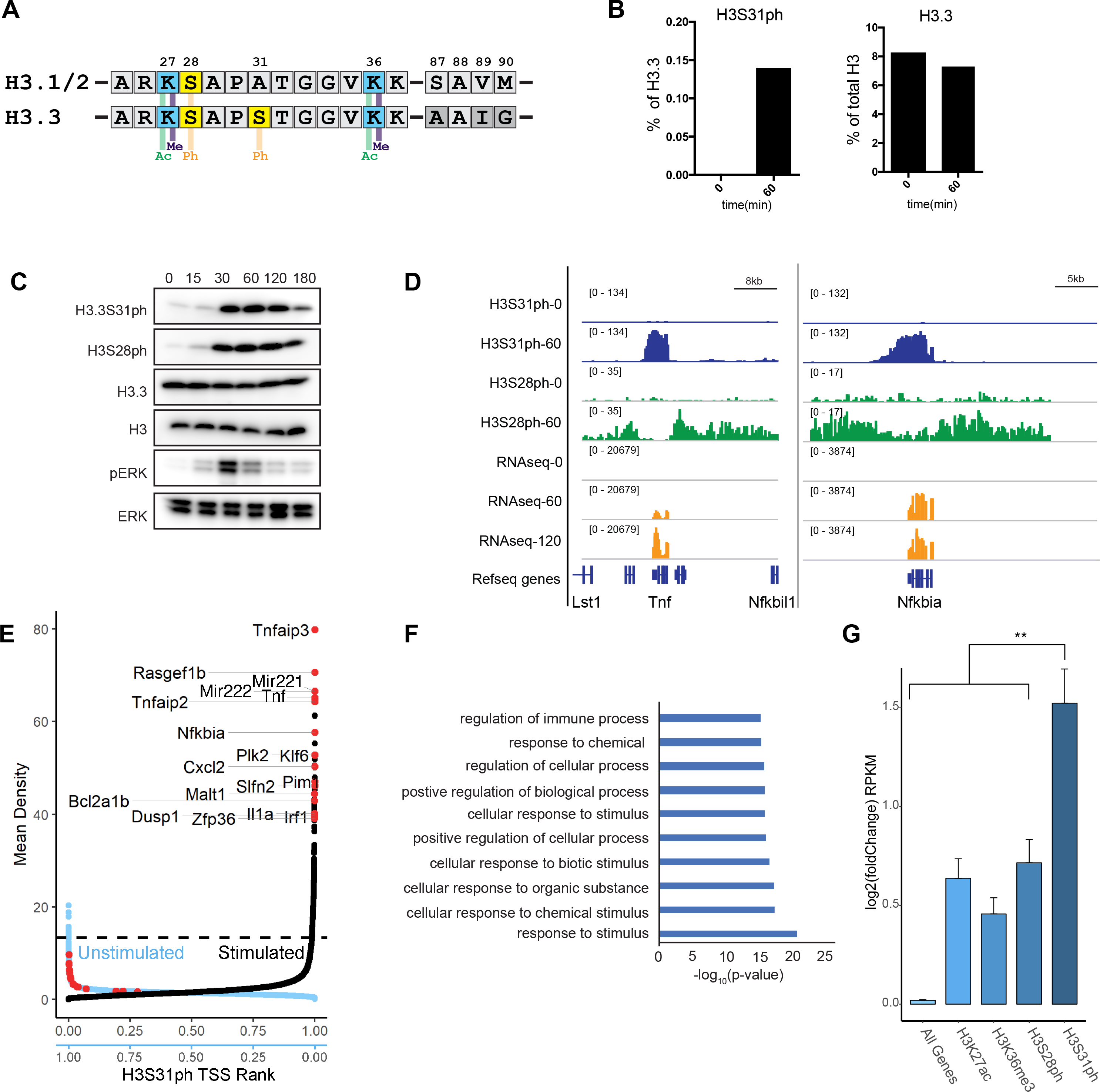
Histone H3 variant, H3.3, is phosphorylated at stimulation induced genes during the macrophage response to pathogen sensing. **(A)** Histone H3 sequence comparison between “canonical” H3.1/2 and variant H3.3 amino-terminal tails (residues 25-37) and the chaperone-specifying motifs in the core domains (87-90), with key histone modifications labeled. **(B)** Quantitative mass spectrometry analysis of phosphorylated H3.3 at Ser 31 (H3.3S31ph), left, and total H3.3 protein, right, in resting (0 minutes) and bacterial lipopolysaccharide (LPS)-stimulated (60 minutes) mouse bone marrow derived macrophages (BMDM). **(C)** Western blot time course analysis of phospho-proteins, H3.3S31ph, H3S28ph, pERK in BMDM response to LPS at increments indicated (in minutes); total H3.3 and Erk as loading controls. **(D)** RNAseq tracks and H3.3S31ph and H3S28ph ChIPseq signals at 0 and 60 minutes of LPS-stimulation at the *Tnf* and *Nfkbia* loci. **(E)** Dual rank order plot of H3.3S31ph ChIP signal density at all genes (TSS-TES) in resting macrophages (ranked in reverse order, right to left, blue X-axis) and stimulated (60’) macrophages (ranked left to right, black X-axis). Dotted line represents the top 1% threshold in stimulated macrophages. Red dots represent top stimulation-induced genes (FDR<0.05, fold change >2 between 0’ and 60’) among the top 0.2% of genes by H3.3S31ph ChIP density and are labeled in the 60’ data. **(F)** Gene ontology (GO) analysis results for the top 1% of genes by rank ordered H3.3S31ph ChIPseq density in LPS-stimulated macrophages (60’). **(G)** Average RNAseq expression (by fold change) for gene sets consisting of all genes and top 1% of genes by ChIPseq density in stimulated macrophages for histone marks H3K27ac (TSS+/−4kb), H3K36me3 (TSS-TES+2kb), H3S28ph(TSS+/− 4kb), and H3S31ph (TSS-TES+2kb). All genes n_all_=16648; Top 1% genes for all ChIP categories n_TOP_=167 (** <0.001). Figure 1B is representative of 2 independent quantitative MS experiments. Figure 1C is representative of 3 or more experiments.

To identify candidate chromatin regulatory mechanisms with a delegated role during cellular stimulation we biochemically purified histones from resting and bacterial lipopolysaccharide (LPS) stimulated macrophages and quantified residue-specific histone post translational modifications (PTMs) by mass spectrometry (MS). Given our interest in the H3.3-specific S31 we targeted peptides containing the H3.3S31 residue in our MS analysis. H3.3S31ph is undetectable in resting macrophages and increases upon stimulation, while the total level of H3.3 protein remains unchanged (Fig. 1B). In support, we developed a specific antibody (fig. S1B-F) and confirmed, by western blot, the stimulation-induced nature and rapid kinetics of H3.3S31ph, paralleling ERK phosphorylation (Fig 1C). Importantly, given the extensive phosphorylation of histones in mitosis, including H3.3S31 (fig.S1B-F) ^18^, the post-mitotic nature of primary mouse bone marrow derived macrophages (BMDM) enabled us to distinguish stimulation-associated histone phosphorylation from mitotic events in bulk populations of cells (fig. S1G).

To establish the genomic location of stimulation-induced H3.3S31ph, we performed chromatin immunoprecipitation followed by whole genome sequencing (ChIP seq) in resting and stimulated (60’ LPS) macrophages. We compared H3.3S31ph localization to H3S28ph. H3S28ph is enriched at promoters, enhancers, and generally across large domains that contain LPS-induced genes, consistent with its role in early events of chromatin activation and transcription^14^. While the H3.3S31ph ChIP signal is enriched in stimulated versus resting macrophages, in striking contrast to H3S28ph, it strictly delineates the “gene bodies” (transcription start site, TSS, to transcription end site, TES) of many LPS-induced genes (Fig. 1D).

A preliminary survey of H3.3S31ph ChIP distribution revealed that its deposition appeared to be specific for stimulation-induced genes (including *Tnf*, *Nfkbia*, *Il1a*, *Il1b*, *Ccl4*, *Cxcl2*, *Tnfaip3*) and is not simply a feature of highly transcribed or constitutively expressed genes (fig. S2). To better evaluate the identity of H3.3S31ph-enriched genes in an unbiased manner and explore the relationship between genic ChIP signal densities of H3.3S31ph and LPS-induced genes, we ranked all annotated genes by H3.3S31ph ChIP signal density (TSS-TES) in resting and stimulated macrophages. This analysis shows that many more genes acquire high-density H3.3S31ph upon stimulation compared with resting cells (Fig. 1E), which is consistent with our MS and other global analysis of H3.3S31ph levels. Additionally, several of the top ranked genes (note, by density, not fold change) are prominent LPS-induced genes, including *Tnfaip3* (A20), *Tnf*, *Il1a*, and *Plk2* (Fig. 1E). We then defined a threshold for the top 1% of genes by H3.3S31ph density in stimulated macrophages (167 genes) for gene ontology analysis and found that the most enriched category is “response to stimulus” (p-value=2.88 × 10^−21^) reflecting the stimulation-induced nature of genes featuring H3.3S31ph (Fig. 1F). We compared the H3.3S31ph chromatin state to other “active” chromatin states including H3K27ac, H3K36me3, and H3S28ph as they relate to stimulation-induced gene expression. Our analysis showed that the top 1% of H3.3S31ph genes (by ChIP density in stimulated macrophages) was highly enriched for stimulation-induced genes (Fig. 1G, fig. S3). Thus, selective deposition of H3.3S31ph at genes with *de novo*, signal-induced transcription indicates a dedicated role in stimulation-responsive transcription rather than constitutive transcription.

In considering possible mechanisms by which H3.3S31ph may regulate transcription, we focused on the gene body localization of this stimulation-dependent histone phosphorylation event. We considered the possibility that H3.3S31ph may be linked to another well-studied histone PTM, the co-transcriptional H3K36me3. H3K36me3 is mediated by a single histone methyltransferase (HMT), SETD2, while members of the NSD family of H3K36-specific methyltransferases can mono- and di-methylate H3K36^19^. SETD2, and specifically the tri-methylation of H3K36, are considered to play an important role in transcription fidelity at highly expressed genes, transcription-associated genic DNA methylation, and mRNA splicing^20–22^. Therefore, we assessed the colocalization and correlation between these two histone PTMs at stimulation-induced genes. We found strikingly similar gene body localization of H3.3S31ph and H3K36me3 in stimulated macrophages, distinct from enhancer and promoter regions delineated by H3K27ac and intergenic regions marked by H3K36me2 (Fig. 2A). Intriguingly, while H3.3S31ph was stimulation-dependent, H3K36me3 was present at modest levels in resting macrophages and increased upon stimulation and induction of associated genes, likely representing a transcriptionally poised state of these genes (Fig. 2A, fig. S3B). While overall, we find enrichment of H3.3 and “active” histone PTMs at LPS-induced genes, H3.3S31ph and H3K36me3 are especially prominent in their enrichment at these genes (Fig. 2A-B, fig. S3A-C). Further, average ChIP density profiling for H3.3S31ph and H3K36me3 across all LPS-induced genes revealed their matching gene-body distribution and stimulation-induced enrichment in this class of genes (Fig. 2A, C). While co-localized at LPS-induced genes, an important distinction between these two histone PTMs is that H3K36me3 is a ubiquitous feature of transcribed genes, while H3.3S31ph appears to have a dedicated function at stimulation-induced genes (fig. S3C).

**Figure 2:**
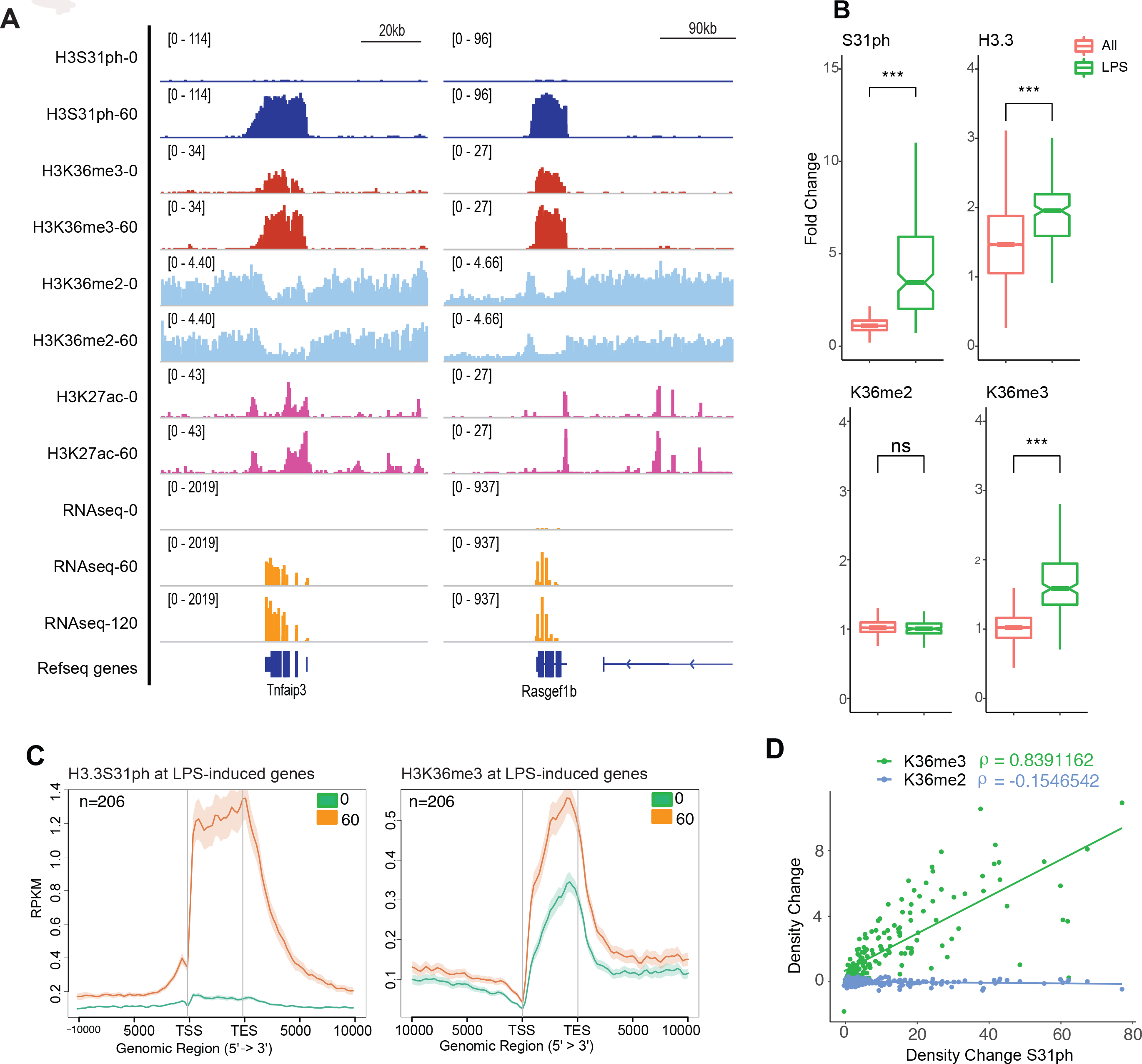
Stimulation-induced H3.3S31ph is deposited in the gene-body of response genes and corresponds with H3K36me3. **(A)** ChIPseq tracks of H3S31ph, H3K36me3, H3K36me3, H3K27ac ChIP (0 and 60 minutes) and RNAseq (0, 60, and 120 minutes) in LPS-stimulated macrophages for Rasgef1b and Tnfaip3. Additional genes and controls are shown in fig. S2. **(B)** ChIP-seq density fold change comparing the set of all genes (All) to RNAseq defined LPS-stimulated genes (LPS) for H3.3S31ph (p=1.22e-96), H3.3 (p=1.63e-25), H3K36me2, and H3K36me3 (p=1.22e-85) by non-parametric Wilcoxon signed-rank test. All genes n_all_=16648; LPS genes n_LPS_=206. **(C)** Average gene profiles of H3.3S31ph (left) and H3K36me3 (right) comparing RNAseq defined LPS-induced genes before and after stimulation. **(D)** Correlation plot showing absolute change (average read density 60’ - 0’ after LPS-stimulation) of H3K36me3 and H3K36me2 association with H3.3S31ph absolute change (average read density 60 - 0).

Thus, H3.3S31ph is a feature of stimulation-responsive chromatin, is rapidly and specifically deposited along the gene bodies of stimulation-induced genes, and at these genes shares a common genomic distribution and stimulated deposition with H3K36me3. These findings suggested cross-talk between these two histone PTMs, and we hypothesized that H3.3S31ph may endow stimulation-induced genes with the capacity for augmented transcription, in part through the stimulation of H3K36me3. To test if H3.3S31ph may determine H3K36me3 densities at LPS-induced genes, we compared H3.3S31ph ChIP density changes between resting and stimulated BMDM with changes in H3K36me3 (catalyzed by SETD2) and H3K36me2 (NSD1, NSD2, NSD3, ASH1L, SMYD2, SETMAR). This analysis demonstrated a high correlation between the density change in H3.3S31ph and H3K36me3 (Spearman’s correlation, 0.8) but not H3K36me2 (Spearman’s correlation, −0.2) (Fig. 2D).

Given this link between H3.3S31ph and H3K36me3 as well as their physical proximity on the H3.3 tail (Fig. 1A), we considered the possibility that H3.3S31ph may directly augment the activity of HMT SETD2, the enzyme catalyzing H3K36me3. To test this hypothesis, we assessed recombinant SETD2-SET domain enzymatic activity *in vitro* on nucleosome substrates assembled from recombinant core histones, either with normal H3.3 tail sequence, or bearing the phospho-mimicking glutamic acid mutation at residue 31 (S31E). Processive SETD2 HMT activity on H3.3K36 was measured by western blot read out during a reaction time course using antibodies specific for K36me2 and K36me3. For comparison, we also performed these assays with the K36me2-specific enzyme NSD2. Under standard assay conditions, both enzymes accumulated their products throughout the 25 minute time course, however, SETD2 activity was potently stimulated by the phospho-mimicking H3.3S31E mutant, while NSD2 activity was substantially reduced (Fig. 3A).

**Figure 3:**
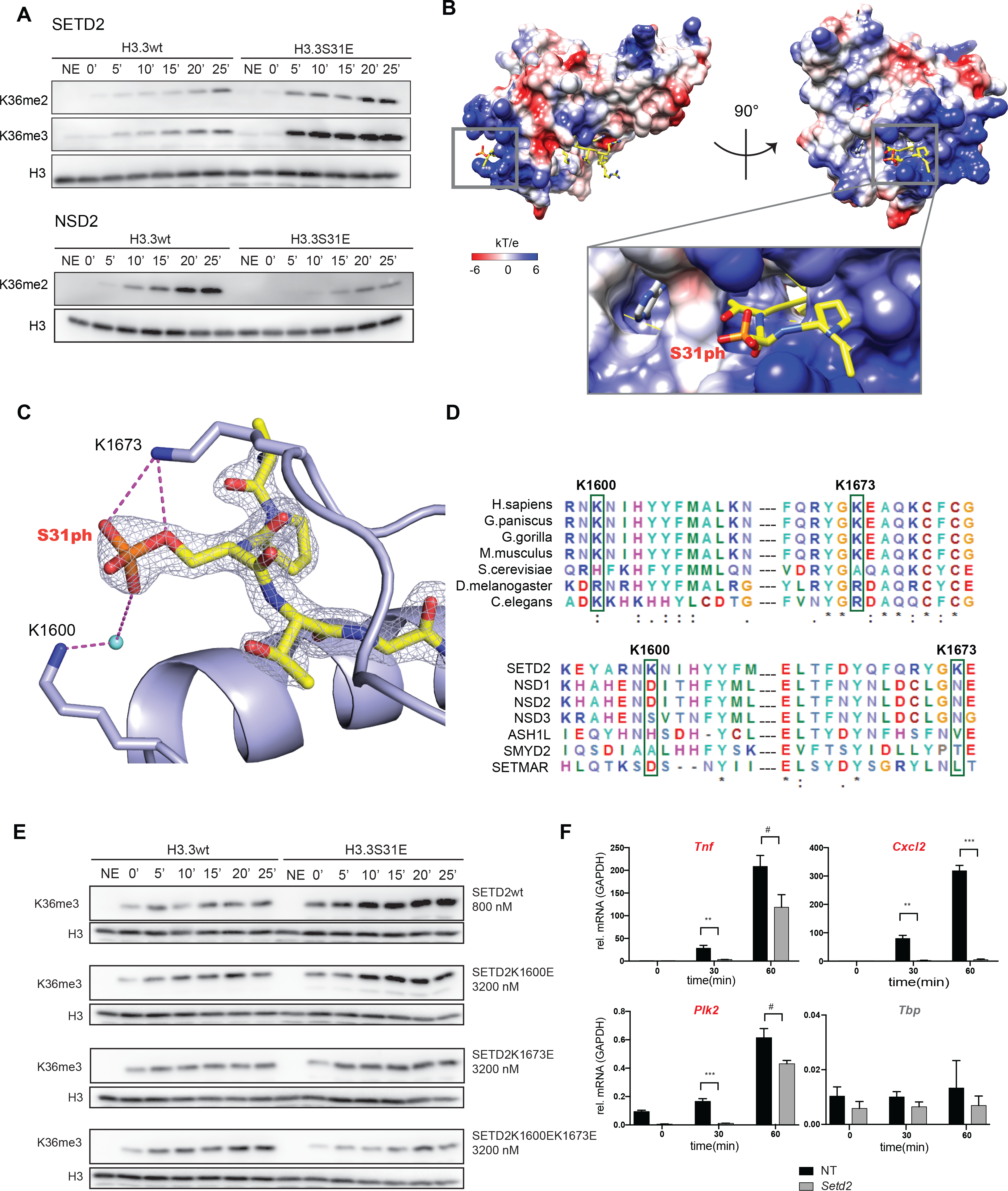
The H3K36me3 methyltransferase SETD2 is stimulated by H3.3S31ph. **(A)** Histone methyltransferase (HMT) assays with SETD2-SET domain and NSD2 full-length enzymes on H3.3wt and H3.3S31E nucleosomes. Reactions were stopped 0, 5, 10, 15, 20 and 25 minutes after adding the enzyme mix to the nucleosomes. Samples were analyzed by Western Blot for H3, HK36me2, and H3K36me3 (NSD2 did not show any signal for H3K36me3). **(B)** Crystal structure of SETD2-H3.3S31phK36M complex. SETD2 is presented as electrostatic potential surface. Electrostatic potential is expressed as a spectrum ranging from −6 kT/e (red) to +6 kT/e (blue). H3.3 peptide is shown as yellow sticks with S31 phosphate group labeled. **(C)** Interaction of the H3.3S31ph phosphate group with K1673 and K1600 of SETD2. The salt bridge bonding and water mediated hydrogen bonding are shown as magenta dashed lines. The peptide is shown as yellow sticks covered by the simulated annealing 2Fo–Fc omit map countered at the 2.0 σ level. The water molecule is shown as aqua blue sphere. **(D)** Top, sequence alignment of SETD2 in different species, highlighting the conserved residues K1600 and K1673, except in S. cerevisiae. Bottom, sequence alignment of different H3K36 methyltransferases highlighting the specificity of residues K1600 and K1673 for SETD2. **(E)** HMT assays with SETD2-SET domain, wild type (wt), K1600E mutant, K1673E mutant and K1600EK1673E double mutant on H3.3wt and H3.3S31E nucleosomes. As the overall activity of the mutant enzymes is reduced, enzyme concentration was titrated to best visualize the ratio of H3.3wt to H3.3S31E activity. **(F)** siRNA knockdown in BMDM for SETD2 or with non-targeting controls (NT) was performed for 2.5 days before LPS stimulation and RT-qPCR for LPS-induced genes *Tnf, Cxcl2, Plk2*, and *Tbp* (constitutively expressed control) at 0, 30, 60 minutes. Setd2 knockdown efficiency is shown in fig. S4A. Enzymatic experiments presented in Figure 3A and 3E were repeated independently, three times, with separate nucleosome and recombinant enzyme preparations and comparable ratios of activities between H3.3wt and H3.3S31E nucleosomes were observed (Figure 3E). siRNA and RT-qPCR experiments (Figure 3F) are representative of 3 independent experiments. **, p<0.01; ***, p<0.001; #, p=0.07, 0.06, for *Tnf*, *Plk2*, respectively, Student t-test.

Structural studies of the SETD2 SET domain have revealed a basic patch along the path of the H3 amino-terminal tail as it extends from the catalytic site^23,24^. We speculated that such a feature could provide the basis of a specific enhanced interaction between SETD2 and H3.3S31ph nucleosome substrates and that these interactions might link the augmented enzymatic activity we observed to structural properties.

Therefore, we solved the crystal structure of the human SETD2 catalytic domain bound to the H3.3 peptide H3.3S31phK36M (S31 phosphorylated, and K36 mutated to M to stabilize the H3.3 peptide in the catalytic site) at 1.78Å (Fig. 3B-C, Table S1). In the resulting structure, the electrostatic surface view of SETD2 shows that the H3.3 peptide is embedded in the substrate-binding channel of SETD2 (Fig. 3B). Notably, the N-terminal fragment of H3.3 extends from the active site to the exit of SETD2 substrate channel, which is exclusively enriched with basic residues. The electron density of the H3.3S31 phosphate group is clearly visualized. Specifically, the hydroxyl oxygen from H3.3S31ph forms a salt bridge with K1673 of SETD2, and water-mediated hydrogen bonding with adjacent K1600 of SETD2 (Fig. 3B-C). Thus, SETD2 K1600 and K1673 provide a channel with positive charge that accommodates and provides charge-complementarity for H3.3S31ph substrate while H3.3K36 is positioned at the active site. Given their potential significance in the observed interactions between H3.3S31ph and SETD2, we evaluated the sequence conservation at and around these lysine residues across phylogeny (Fig. 3D top) and within H3K36 methyltransferases (Fig. 3D bottom). Remarkably, we find that the basic residues K1600 and K1673 are highly conserved in metazoan SETD2 (conserved across vertebrates and replaced by highly similar Arg in *C. elegans* and *D. melanogaster* and His in *S. cerevisiae*). In contrast to this high degree of cross-species conservation within SETD2 orthologs, other H3K36 HMTs (NSD family, etc.) frequently replace these basic residues with acidic or polar amino acids (Fig. 3D, bottom).

To directly assess the function of these conserved SETD2 lysine residues that engage in specific interactions with H3.3S31ph, we generated recombinant SETD2 SET-domain proteins with mutated lysines, individually and combined (K1600E, K1673E, and K1600E/K1673E). Wild type and mutant SETD2 enzymes were then assessed for their activity on unmodified as well as H3.3S31E-containing nucleosomes. As before, we observed potent stimulatory activity of H3.3S31E nucleosomes over unmodified nucleosomes (Fig. 3A, E), however, H3.3S31E-augmented SETD2 activity was decreased in single mutants (K1600E and K1673E) and reversed in the double K1600E/K1673E SETD2 mutant.

Together, our cellular, epigenomic, and structure-function studies suggest H3.3S31ph-augmented SETD2 activity as a feature of enhanced stimulation-induced transcription. This indicates a mechanism by which stimulation-induced genes may be endowed with preferential access to (and dependency on) SETD2 for rapid, high-level expression. To test this hypothesis, we performed SETD2 siRNA knockdown in BMDM before LPS stimulation. We find that expression of LPS-induced genes with H3.3S31ph, *Tnf*, *Plk2*, *Cxcl2*, is highly dependent on SETD2, compared with constitutively expressed *Tbp* (Fig. 3F, fig. S4A).

Functional perturbations of histone genes are made difficult by their essential role in diverse cellular function and their genetic complexity (there are 15 copies of the H3 gene in mouse and human). However, because there are only two genes for the histone H3.3 variant containing the S31 residue (*H3f3a* and *H3f3b*) we were able to target these genes by CRISPR. H3.3 is required for embryogenesis^25–28^ and spermatogenesis^29^. Further, as we find here (fig. S3), H3.3 is enriched at inflammatory genes, though its function in this context is unknown^6,30^. To study the function of H3.3 in inflammatory gene induction we generated *H3f3a/H3f3b* double knockout (DKO) RAW264.7 (macrophage-like) mouse cell lines through CRISPR targeting of both *H3f3a* and *H3f3b*. Given its critical role in development, we also selected a hypomorphic (HYPO) RAW264.7 clone, with a null *H3f3a* allele and hypomorphic *H3f3b* allele (fig. S4B).

These wild type, DKO, and HYPO macrophage cell lines were then assessed for their ability to induce inflammatory genes following stimulation with LPS in the absence of the H3 protein containing the H3.3-specific S31 residue. While these cell lines grow comparably (not shown), assessment of their ability to rapidly respond to stimulation by RNAseq revealed substantial decreases in induced expression of LPS-induced genes in both DKO and HYPO macrophage cell lines (Fig 4A-B, fig. S4C-D). At 60 minutes and 120 minutes following stimulation with LPS, we observed a global reduction in LPS-induced gene expression in DKO and HYPO cell lines (Fig. 4B-C, fig. S4D). We found that LPS-induced genes characterized by the highest levels of H3.3S31ph were expressed, on average, at 3-times the level of all LPS-induced genes and also had consistently decreased expression in H3.3 HYPO and DKO cells (Fig. 4C-D, fig. S4D).

**Figure 4:**
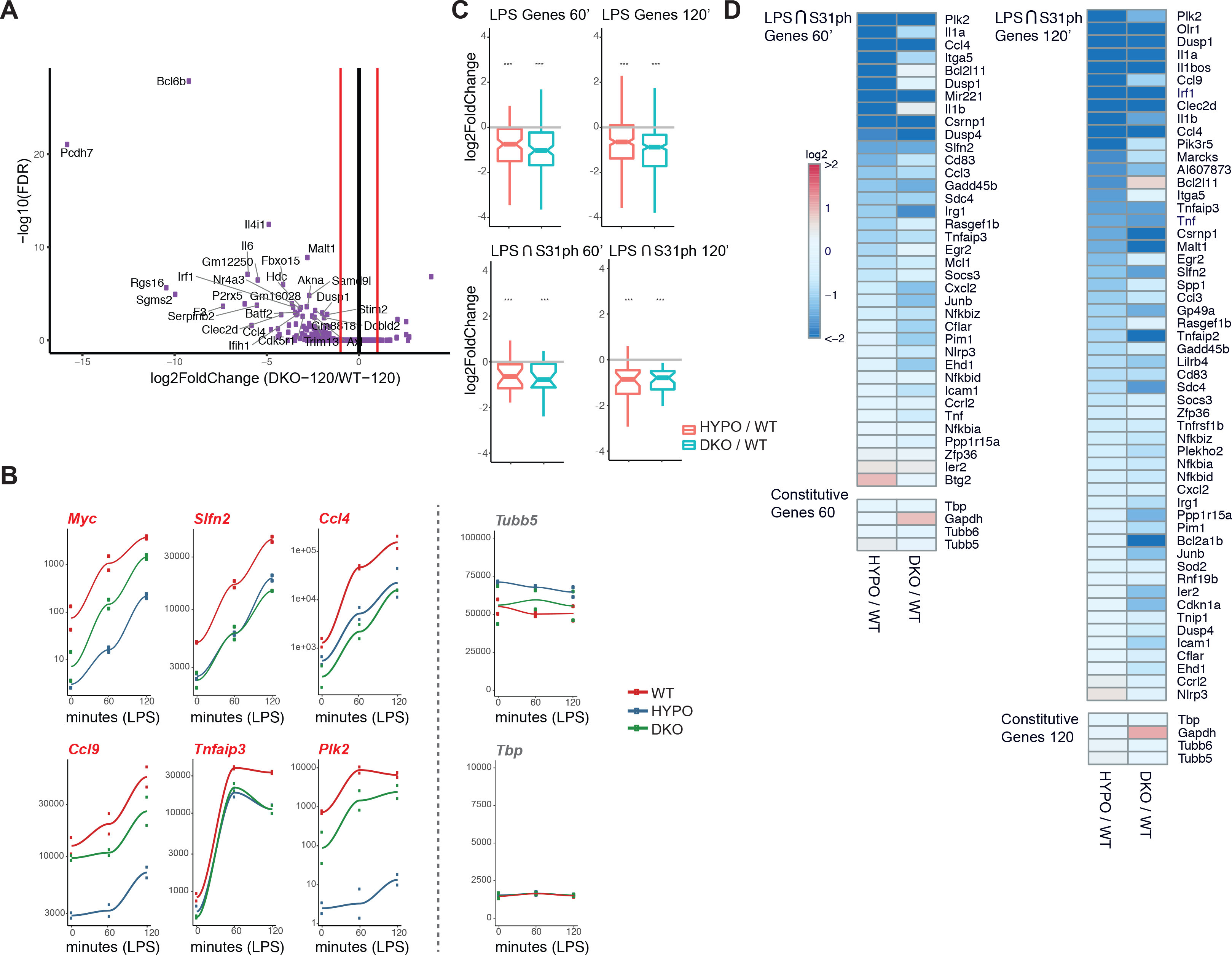
H3.3 is critical for stimulation-induced transcription of inflammatory genes. **(A)** RNAseq scatter (“volcano”)-plot analysis, log2 fold-change and −log10(FDR), of DKO compared to wt RAW247.6 at 120 minutes. **(B)** Time course plots of mean RNAseq expression (RPKM) from two experiments at time points 0’, 60’, and 120’ after LPS-stimulation for experiments performed in wild-type (WT), hypomorph (HYPO), and double-knockout (DKO) RAW247.6 cell lines at LPS-induced genes *Myc, Ccl9, Slfn2, Tnfaip3, Ccl4, Plk2*, and constitutively expressed genes *Tubb5 and Tbp*. **(C)** Ratio of RNAseq fold change (log_2_) for HYPO or DKO compared with WT at 60’ and 120’ LPS stimulation for all LPS induced genes (top) and for the intersection of top H3.3S31ph genes and LPS induced genes (bottom). ***<0.0001 by lower-tailed one-sample t-test (distribution below zero). **(D)** Heat map of fold change (log_2_) for top H3.3S31ph genes among LPS-induced genes (left, 60 minutes; right, 120 minutes) with control constitutively expressed genes below. RNAseq was performed with two biological replicates per condition.

The inflammatory gene induction defect in H3.3 HYPO and DKO cells responding to LPS occurs despite the presence of 13 other copies of H3.1/2, abundantly expressed in these rapidly cycling cells. Given that the only H3.3 “tail” sequence difference is S31 and our results that demonstrate a dedicated role for H3.3S31ph at stimulation responsive genes, we suggest that H3.3S31ph contributes to the function of H3.3 that we observe in these experiments (Fig. 4). Our structural and enzymatic studies of Setd2 activity (Fig. 3) highlight specific biophysical mechanisms that may link H3.3 to augmented transcription. However, key unknowns remain on the function of this ancestral H3.3 variant, including the relative function of the H3.3S31 residue and its phosphorylation, the signaling pathways that link stimulation to H3.3S31ph in chromatin, and the breadth of the mechanisms that we describe here, both across species and cell types.

Dedicated mechanisms enabling rapid stimulation-induced transcription are relevant to diverse cell responses and disease states, and may represent more selective therapeutic targets than the general transcription machinery^31–33^. In the context of inflammatory gene induction, numerous studies have revealed signals, TFs, and chromatin features that drive stimulation responsive genes (reviewed in ^2,34,35^). However, explanation of inducible genes’ preferential access to the transcription apparatus and suitability for speed and scope of transcription in the form of dedicated chromatin mechanisms have remained obscure. Our epigenomic and biochemical studies link selectively deposited H3.3S31ph at stimulation-induced genes to augmented SETD2 activity and co-transcriptional H3K36me3, enabling rapid and high-level transcription of these genes. Together with our previous characterization of H3S28 phosphorylation in early stimulation-induced chromatin activation^14^, these studies reveal mechanisms for the dedicated role of histone phosphorylation in *de novo* transcription. We propose that selectively employed deposition of histone PTMs at these genes, including H3.3-specific H3.3S31ph, provides a signature that specifies preferential access to the transcription apparatus, endowing cells with the essential capacity for rapid and selective environmental responsiveness.

## Acknowledgements

This work was supported by the following funding sources: R01GM040922 (C.D.A), R00GM113019 (S.Z.J), CIPSM (S.B.H.). We thank John Zinder for contributing the SETD2-pETduet-smt3 construct; Congcong Lu, Simone Sidoli (lab of B.A.G.) for H3.3 peptide analysis; members of Weill Cornell Applied Bioinformatics Core, Doron Betel, Paul Zumbo, Friederike Dundar, and Luce Skrabanek for suggestions and assistance with bioinformatics; Alexey Soshnev for help with figures.

## Author Contributions

A.A. and S.Z.J. designed the study, performed biochemical, cellular, and epigenomic experiments and analyzed the data. S.Y. performed structural studies supervised by H.L.. L.E.R., C.D., A.W.D., J.Q.J., A.L.M., A.R., performed experiments and analyzed data supervised by S.Z.J.. T.P. assisted A.A. with nucleosome assembly and enzymatic assays. T.A. and S.B.H. developed and tested the H3.3 antibody. S.L. performed mass spectrometry studies supervised by B.A.G.. S.Z.J. wrote the manuscript with input from all authors. C.D.A., H.L., and S.Z.J. supervised the study.

## Materials and Correspondence

S.Z.J.

---

**Figure S1:**
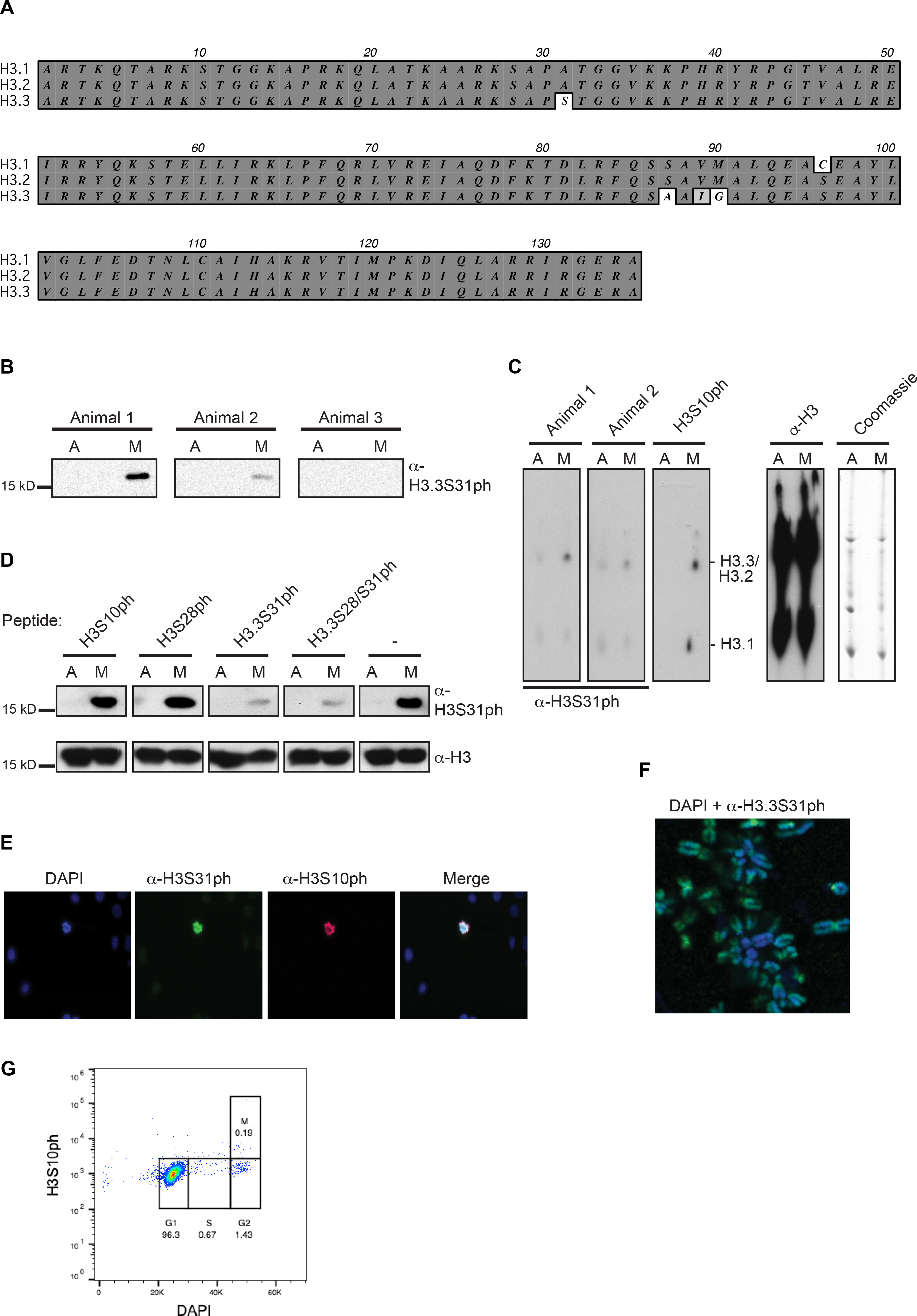
Determination of anti-H3.3S31ph antibody specificity. **(A)** Alignment showing H3.1, H3.2 and H3.3 and the differing amino acids in core and tail. **(B)** Immunoblot with acid-extracted histones from asynchronous growing “A” or nocodazole-arrested mitotic “M” HeLa cells using bleeds from three rabbits immunized with H3.3S31ph peptides. Bleeds from animals 1 and 2 show a signal of the molecular weight of histone H3 only with mitotic samples. **(C)** Immunoblot with acid-extracted histones from asynchronous “A” or mitotic “M” histones separated by 2D triton-acid urea (TAU) gels (left) that allow a separation of histone variants due to charge and amino acid differences. The bleed from animal 1 shows a signal of the size of H3.3. Coomassie blue staining of the gel and staining of the membrane with anti-H3 served as loading control (right). **(D)** Peptide competition experiment to determine antibody-specificity. Asynchronous “A” or mitotic “M” histones were separated by SDS-PAGE gels and blotted onto PVDF membranes. H3.3S31ph antibody from animal 1 was pre-incubated with diverse peptides or without any peptide, as indicated, before adding it to the PVDF membrane. Staining with anti-H3 antibody shows equal loading. **(E)** Deconvolved immunofluorescence microscopy images of asynchronously growing HeLa cells co-stained with DAPI (DNA, blue), anti-H3.3S31ph (animal 1, green) and anti-H3S10ph (marker of mitotic cells, red). Merged picture is shown on the right. Note that only mitotic cells, as apparent from stronger DAPI-staining and apparent H3S10ph signal, are H3.3S31ph positive. **(F)** Deconvolved image of chromosome spread from mitotic HeLa cells co-stained with DAPI (blue) and anti-H3.3S31ph (animal 1, green). Notice the stronger staining of H3.3S31ph at peri-centromeric regions, as has been shown previously. **(G)** Cell cycle analysis of BMDMs by FACS using DAPI and H3S10ph, with mitotic index gate shown, indicating post-mitotic nature of BMDMs.

**Figure S2:**
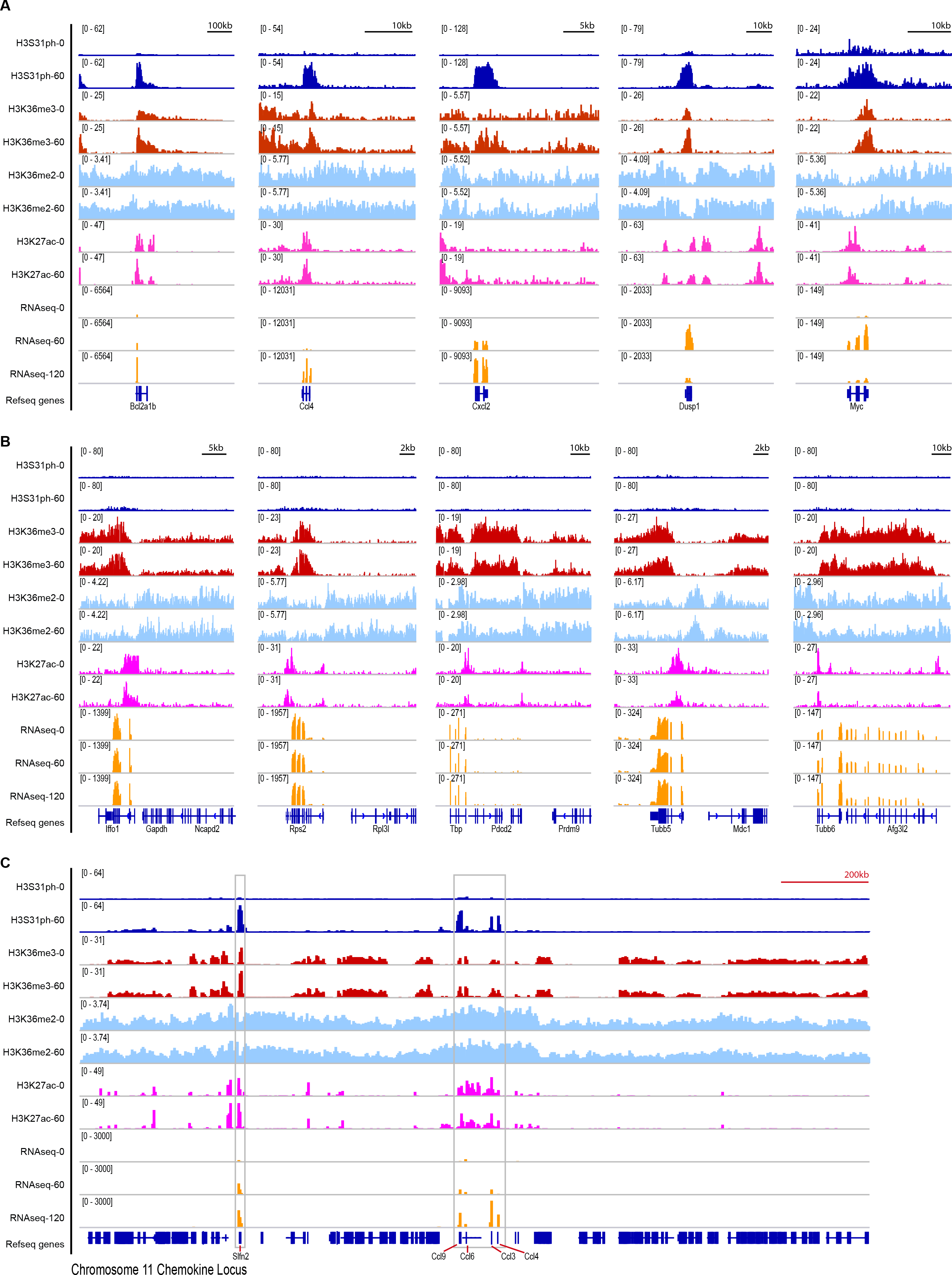
H3.3S31ph is deposited in the gene-body of response genes but not constitutively expressed genes. Additional examples of H3 PTMs including H3.3S31ph (as in Figure 2A) at **(A)** LPS-induced genes *Bcl2a1b, Ccl4, Cxcl2, Dusp1, Myc*; **(B)** constitutively expressed (“housekeeping”) genes *Gapdh, Rps2, Tbp, Tubb5, Tubb6*; **(C)** across the gene dense chromosome 11 chemokine locus (>1Mb) containing LPS-induced genes *Slfn2, Ccl9, Ccl6, Ccl3, Ccl4*.

**Figure S3:**
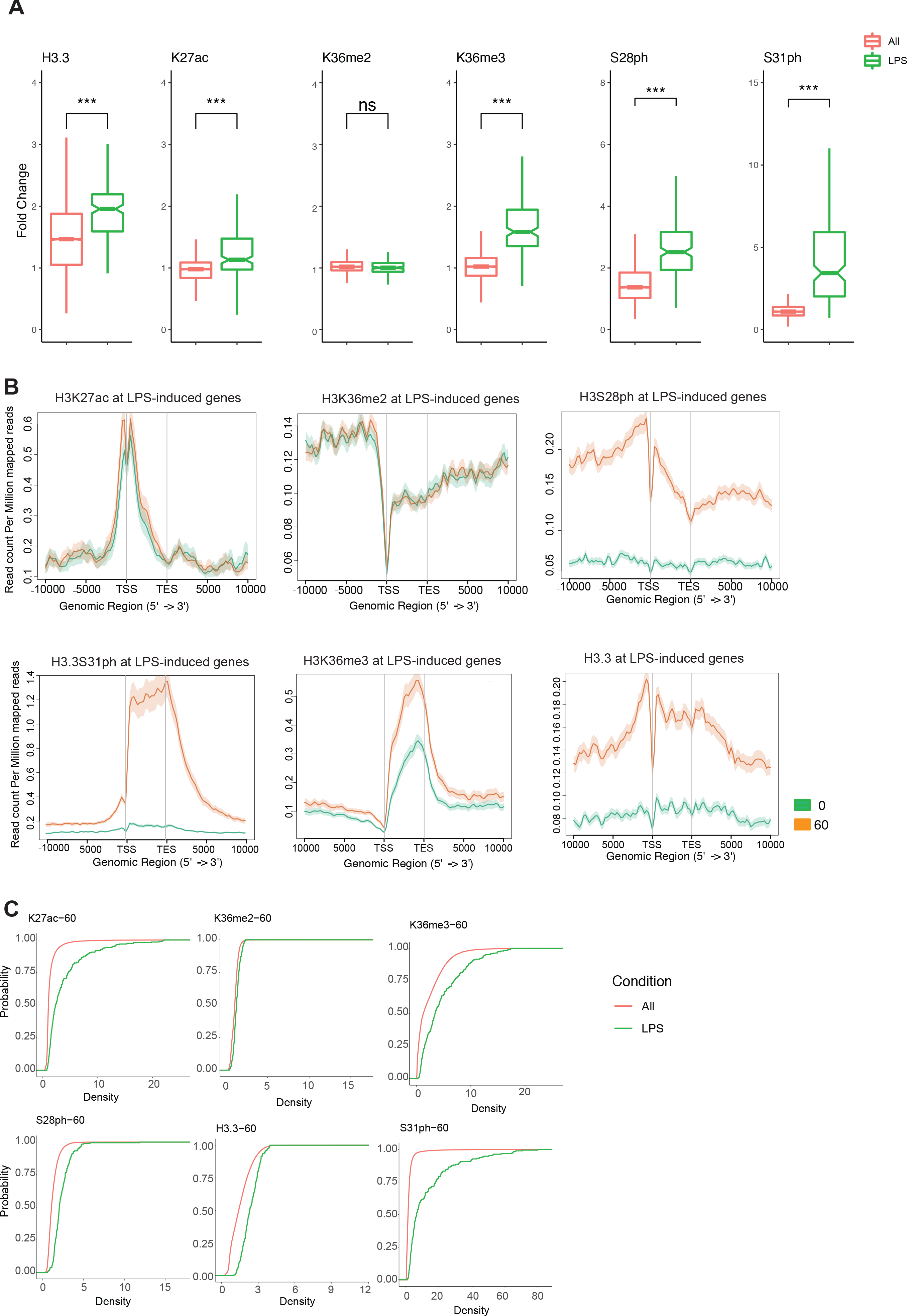
H3.3S31ph and other H3 PTMs at response genes and after stimulation. **(A)** Additional examples (H3K27ac and H3S28ph) of ChIP-seq density fold change comparing the set of all genes (All) to RNAseq defined LPS-stimulated genes to RNAseq defined LPS-stimulated genes (LPS) as shown in Figure 2B for (H3.3, K36me2, K36me3, S31ph) **(B)** Average gene profiles (in addition to H3.3S31ph and H3K36me3 in Figure 2, shown here are H3K27ac H3K36me2, H3S28ph and H3.3) comparing RNAseq defined LPS-induced genes before and after stimulation. **(C)** Cumulative distribution function (CDF) plots for H3K27ac H3K36me2, H3K36me3, H3S28ph, H3.3 and H3.3S31ph reveal selective role of H3.3S31ph compared with ubiquitous role of H3K36me3. ***<0.0001 by student t-test.

**Figure S4:**
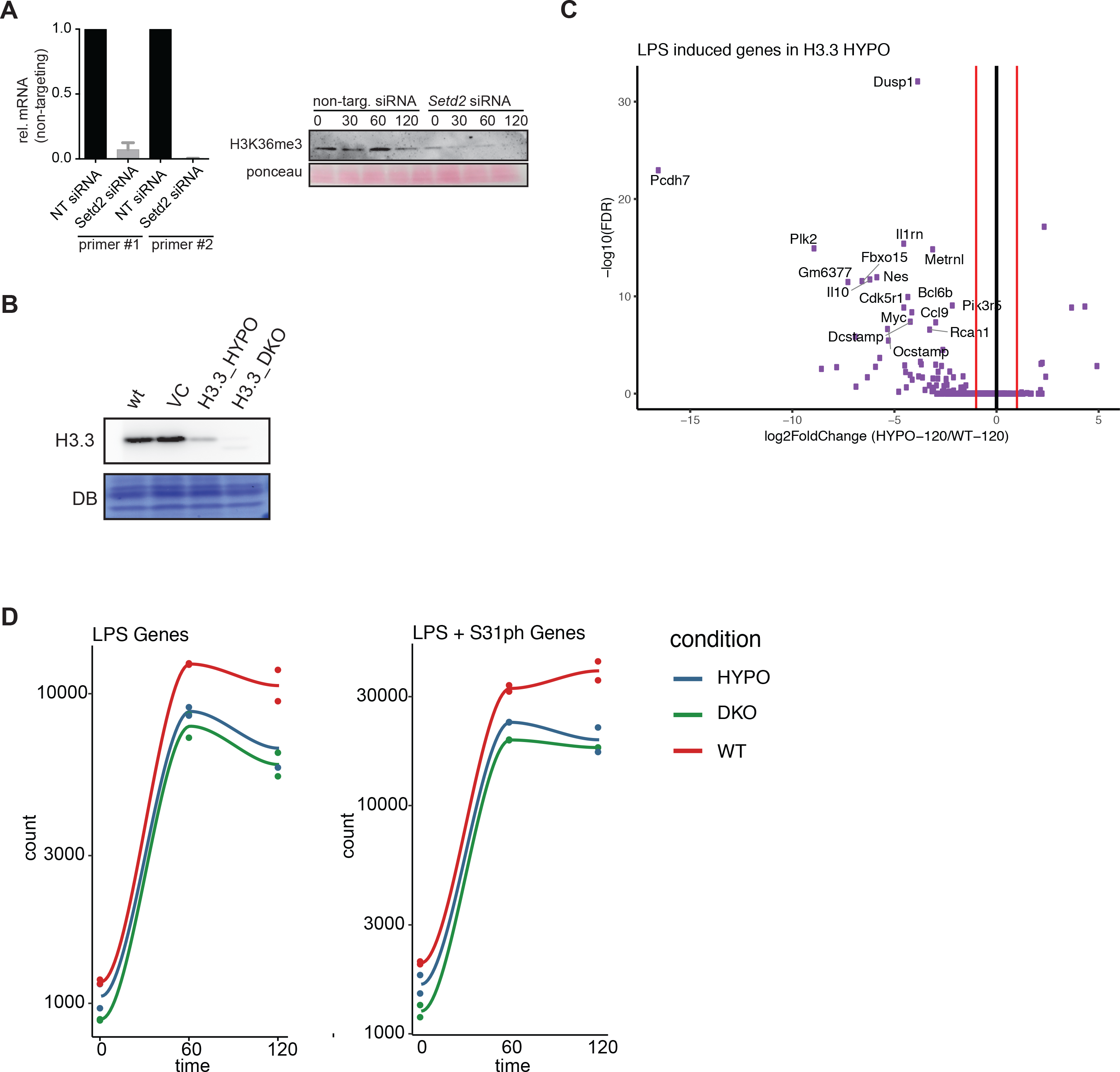
Characterization of Setd2 knock down and H3.3 mutant RAW247.6 cell lines. **(A)** siRNA knockdown validation of siRNA for *Setd2* using two RT-PCR primers (left) and western blot for H3K36me3 as a surrogate of Setd2 activity (right). **(B)** Western Blot for H3.3 comparing wild-type (WT), vector control (VC), hypomorph (HYPO), and double-knockout (DKO) RAW247.6 cell lines, membrane was stained with direct blue (DB) for equal loading. **(C)** RNAseq scatter (“volcano”)-plot analysis, log2 fold-change and −log10(FDR), of HYPO compared to WT RAW247.6 at 120 minutes. Matches Fig. 4A, DKO plot. **(D)** Time course plots of mean RNAseq expression (RPKM) from two experiments at time points 0’, 60’, and 120’ after LPS-stimulation for experiments performed in wild-type (WT), hypomorph (HYPO), and double-knockout (DKO) RAW247.6 cell lines at LPS-induced genes and at top H3.3S31ph genes among LPS-induced genes.

**Table S1.**
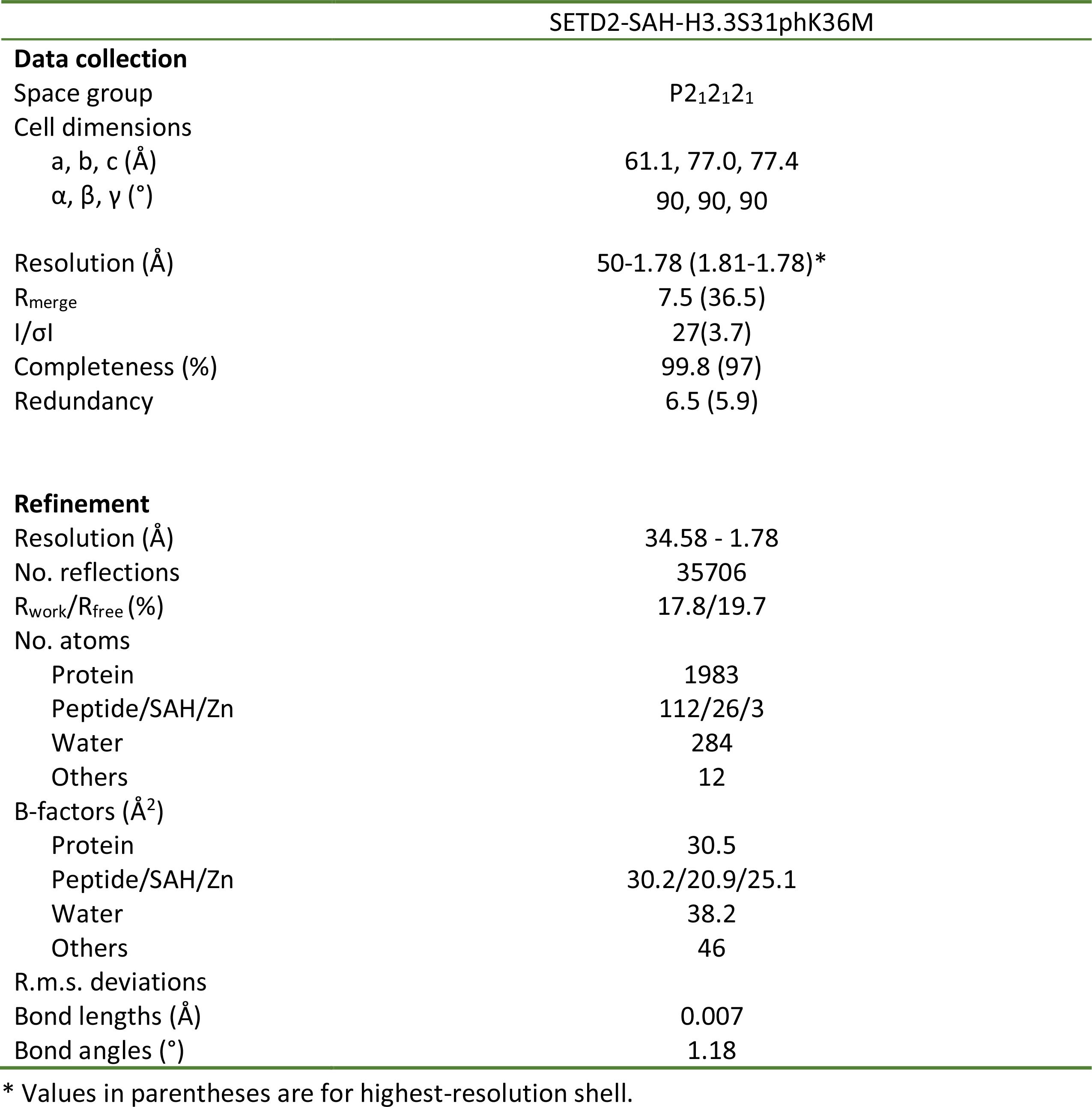
Data collection and refinement statistics

## Materials and Methods

### ChIP-seq data processing and analysis

H3S31ph, H3K36me3, H3K36me2, H3K27ac, H3S28ph, and H3.3 ChIP-seq analyses were performed in bone marrow derived macrophages (BMDM) with an average range of 20-25 × 10^6^ reads per independent ChIP-seq experiment. ChIP-seq reads were mapped to the mm10 genome using Bowtie2 v.2.3.4.1^1^ with the following parameters: -p 8 ‒k 1 ‒N 1. The aligned reads underwent three stages of filtering using SAMtools v.1.5^2^. First, the unmapped, non-primary, qc failed, and multi-mapped reads were discarded. PCR duplicates were then marked by Picard Tools v.2.14.0 (http://broadinstitute.github.io/picard/) using ‘VALIDATION_STRINGENCY=SILENT and REMOVE_DUPLICATES=false” options and removed by SAMtools (−F 1796). Then, chromosome M and scaffolds were removed to create the final filtered bam file. The final bam files were used to generate average profiles for RNA-seq define LPS-stimulated genes at time 60 for H3S31ph signal using ngs.plot v.2.61^3^ at genebody using the following parameters: −FL 200 −MW 2. For visualization in IGV v.2.3.94^4^, the final bam files were converted to a tiled data file (.tdf) using igvtools v.2.3.98^5^ including duplicates. Final bam files were converted to bigWig files of read coverages normalized to 1x depth of coverage as reads per genomic content (RPGC) using deeptools v2.5.4^6^ bamCoverage. To obtain a tab-delimited file of average scores comprised of all bigWig files for each experiment, deeptools multiBigwigSummary performed the analysis for regions defined by a General Transfer Format (GTF) vM3 Annotation BED file. The BED file was constructed using the BEDOPS v.2.4.29^7^ gtf2bed conversion utility and, depending on strand direction, extending the feature at both the start and end position by 2kb (H3S31ph, H3K36me3, H3K36me2, H3.3) or 4kb (H3S28ph, H3K27ac) to account for promoters (+/−2kb) or histone marks found outside of gene body (+/− 4kb). The resulting tab-delimited file of read densities was used for downstream analysis in R v.3.4.0^8^. Top H3S31ph genes were defined by a 2-fold or greater increase in H3S31ph enrichment at time 60 after LPS stimulation with FDR < 0.05. Top genes for all other epigenetic marks, such as H3K27ac, H3K36me3, H3S28ph, were defined in the same manner. The top H3S31ph genes enriched at time 60 were used as a target list for gene ontology analysis by the tools Gorilla^9^ and REViGO^10^.

### RNA-seq data processing and DESeq2 analysis

Paired-end RNA-seq reads were obtained from biological triplicates at times 0, 60, and 120 after LPS stimulation in BMDMs. Single-end RNA-seq reads were also obtained from technical duplicates at times 0, 60, and 120 after LPS stimulation for KO comparisons for WT BMDM, cell line hypomorph 3.205, and cell line knockout 264. Both paired-end and single-end RNA-seq were processed the same. The fastq files underwent adapter trimming and quality control analysis using wrapper Trim Galore v.0.5.0. The resulting trimmed fastq files were aligned to the GENCODE vM3 transcriptome in mm10 using STAR aligner v.2.4.2^11^ with default settings. The utility featureCounts^12^ from Subread v.1.4.6 was used to calculate raw counts reads per gene to be used as input for differential expression analysis by DESeq2^13^.

### Antibodies

a-H3.3S31ph (developed by Pineda Antikörper-Service), a-H3S28ph (clone E191, ab32388 Abcam), H3.3 (09-838, EMD), a-p44/42 MAPK, Erk1/2 (4695 Cell Signaling), a-phospho-p44/42 MAPK (Erk1/2) (4370, Cell Signaling), a-H3 (ab1791 Abcam), a-H3K27ac (39133, Active Motif), a-H3K36me3 (61021, Active Motif), a-H3K36me2 (2901, Cell Signaling).

### a-H3.3S31ph Antibody Development

For the generation of an H3.3S31ph-specific polyclonal antibody, a peptide spanning amino acids 26 to 37 from H3.3 containing phosphorylated serine 31 (RKSAPS(ph)TGGYKK, note the exchange of V35Y due to enhanced immunicity) was used for immunization of three rabbits by the Pineda - Antikörper-Service company (Berlin, Germany). Last bleed from animal 1 was affinity purified and used in this study. Antibody specificity was tested in immunoblots and 2D-Triton Acid Urea (2D-TAU) gels with acid-extracted histones as described previously^14^. Peptide competition experiments were done as described previously^15^ using peptides that were N-terminally biotinylated and synthesized with higher than 80% purity by GenScript USA Inc. All peptides contained the general H3.3 sequence (aa 20-39; BIO-LATKAARKSAPSTGGVKKPH) with respective phosphorylations on serines 10, 28 and/or 31. For Immunofluorescence microscopy HeLa Kyoto cells were grown on coverslips, washed, fixed, permeabilized and stained as descibed previously^16^. Chromosome spreads were generated as described^17^. Wide-field fluorescence imaging was performed on a PersonalDV microscope system (Applied Precision) equipped with a 60x/1.42 PlanApo oil objective (Olympus), CoolSNAP ES2 interline CCD camera (Photometrics); Xenon illumination and appropriate filtersets. Iterative 3D deconvolution of image z-stacks was performed with the SoftWoRx 3.7 imaging software package (Applied Precision).

### Chromatin Immunoprecipitation

As previously described in Josefowicz et al., 2016.

### Primary Cell Culture

As previously described in Josefowicz et al., 2016. HeLa Kyoto cells were grown as described^15^.

### Cell Culture, siRNA transfection

For siRNA transfection RAW cells were reverse transfected with Lipofectamine RNAiMAX (Life Technologies) and ON-TARGETplus SMARTpool siRNAs against mouse SETD2, CHK1 and CHK2. After 72h, cells were either harvested for gene expression or western blot analysis.

### RNA extraction, quantitative real-time PCR and RNA sequencing

RNA was isolated using RNAeasy Kit (Quiagen). For RT-PCR extracted RNA was treated with DNAse and cDNA was synthesized using High-Capacity cDNA Reverse Transcription Kit (Applied Biosystems). qPCR was performed using SYBR green dye (Applied Biosystems) and normalized to GAPDH. For RNA sequencing libraries were prepared using according to the Illumina TruSeq protocol and were sequenced on Illumina HiSeq 2500 / NextSeq 500.

### Antibody-based methods

(flow cytometry and western blotting) As previously described in Josefowicz et al., 2016.

### Mass Spectrometry Analysis of Histone Post-Translational Modifications

As previously described in Josefowicz et al., 2016.

### Nucleosome reconstitution

All histones were expressed and purified as previously described (Ruthenburg et al., 2011). Nucleosome Assembly Octamers were reconstituted as described (Ruthenburg et al., 2011). The 601 nucleosome positioning sequence was used for nucleosome reconstitution (Lowary and Widom, 1998). The DNA was amplified by PCR using HPLC purified primers containing a biotin tag on the 5’ end to produce 189 bp linear DNA and purified using QIAEXII kit (Qiagen). Nucleosomes were assembled using the standard step-wise dialysis method (Dyer et al., 2004).

### Bacterial recombinant protein

Human SETD2_1347-1711_ (original plasmid was a generous gift of Danny Reinberg) and point mutants were cloned into pETduet–smt3 (Mossessova E, Lima CD, 2000). The SETD2 wt and mutant fragments were expressed with an N-terminal His-tag in Rosetta (DE3, pLysS) cells with LB Media for 18 h at 17°C by induction with 0.5 mM Isopropyl β-D-1-thiogalactopyranoside (IPTG). E. coli cells were resuspended in50 mM Tris pH 8.0, 500 mM NaCl, 1 mM PMSF, 2 mM BME, 10% glycerol, 10 mM imidazole supplemented with ROCHE COMPLETE protease inhibitors. After lysis with tip sonicator and centrifugation the cleared lysate was incubated for 1h with Ni-NTA resin slurry (Clonetech). After washing beads with the same buffer, the protein was eluted. The samples were incubated with Ubiquitin-like protease (Ulp) overnight at 4°C and subsequently incubated again with Ni-NTA resin to remove protease and cleaved tag. Supernatant was further purified by size-exclusion chromatography (Superdex 75, GE Healthcare).

### HMT assay

Standard HMT assays were performed in a total volume of 20 μL containing HMT buffer (50 mM Tris-HCl, pH 8.5, 50mM NaCl, 5 mM MgCl_2_, and 1 mM DTT) with 100 uM S-Adenosylmethionine (NEB) and 1.2ug of nucleosomes. The enzymes used were 30nM NSD2 full-length (Reaction Biology Corp), 800 nM SETD2-SET wt, and 3200 nM of SETD2K1600E, SETD2K1673E, SETD2K1600EK1673E. The reaction mixtures were incubated for 0,5,10,15,20 and 25 min at 30°C and stopped by adding 20ul of Laemmli Buffer. The results were analyzed by Western Blot.

### Crystallography study of SETD2-H3.3S31phK36M complex

Human SETD2 catalytic domain (residues 1434–1711) was expressed in E. *coli* and purified as previously described (Yang et al. 2016). Crystallization was performed via vapor diffusion method under 277K by mixing equal volumes (0.5ul) of SETD2-H3.3_29-42_S31phK36M-SAM (1:5:10 molar ratio, 8mg/ml) and reservoir solution containing 0.2M potassium thiocyanate, 0.1M Bis-Tris propane, pH 8.5, and 20% PEG 3350. The crystals were briefly soaked in a cryo- protectant drop composed of the reservoir solution supplemented with 20% glycerol and then flash frozen in liquid nitrogen for data collection. Diffraction data were collected at Shanghai Synchrotron Radiation Facility beamline BL17U under cryo conditions and processed with the HKL2000 software packages. The structures were solved by molecular replacement using the MolRep program (Vagin and Teplyakov 2010), with the SETD2-H3.3K36M complex structure (PDB code: 5JJY) as the search model. All structures were refined using PHENIX (Adams et al. 2010) with iterative manual model building with COOT (Emsley and Cowtan 2004). Detailed structural refinement statistics are in Supplemental Table S1. Structural figures were created using the PYMOL (http://www.pymol.org/) or Chimera (http://www.cgl.ucsf.edu/chimera) programs.

### *In vitro* kinase assay and dot blot

Recombinant CHK1 kinase (Sigma) was incubated with kinase buffer (40mM HEPES pH7.4, 20mM MgCl2), Magnesium/ATP cocktail (EMD) and histone tail peptides for overnight at 37°C (Total reaction 15ul, 2ug Peptide, Mg(4.5mM)/ATP(30uM) cocktail and 4ng Enzyme). The samples were then added with 5ul of 0.5%SDS followed by boiling for 5min at 95°C. The samples were dropped on a dry nitrocellulose membrane and probed with a-H3S31ph antibody.

### CRISPR targeting of H3.3

CRISPR targeting *H3f3b* and *H3f3a* was performed in RAW264.7 cells using methods described in Ran et al. 2013^18^. Targeting was done consecutively first targeting *H3f3b*, then using *H3f3b* mutants to target *H3f3a*.

The gRNAs (Primers caccTAGAAATACCTGTAACGATG forward aaacCATCGTTACAGGTATTTCTA reverse for *H3f3a* and caccGAAAGCCCCCCGCAAACAGC forward aaacGCTGTTTGCGGGGGGCTTTC reverse for *H3f3b*) were cloned into PX458 (from Addgene) and sorted for GFP 24h after transfection, cells were first sorted as bulk and after recovery sorted into single cell clones. Positive clones were tested by PCR, sequencing and Western Blot.

